# Arabidopsis root apical meristem adaptation to an osmotic gradient condition: an integrated approach from cell expansion to gene expression

**DOI:** 10.1101/2024.07.19.604290

**Authors:** Selene Píriz Pezzutto, Mauro Martínez-Moré, María Martha Sainz, Omar Borsani, Mariana Sotelo Silveira

**Affiliations:** Laboratorio de Bioquímica, Departamento de Biología Vegetal, Facultad de Agronomía, Universidad de la República, Avenida Garzón 780, Montevideo, CP 12900, Uruguay

**Keywords:** osmotic stress, gradient, root growth, Arabidopsis, *TTL1*

## Abstract

Climate change triggers abiotic stress, such as drought and high salinity, that can cause osmotic stress. Water availability can limit plant growth, and the root tip tissues initially sense it. Most experiments destined to understand root growth adaptation to osmotic stress apply homogeneous high osmotic potentials (osmotic shock) to shoots and roots. However, this treatment does not represent natural field conditions where a root may encounter increasing osmotic potentials while exploring the soil. Osmotic shock severely reduces root growth rate, decreasing cell division in the proximal meristem and reducing mature cell length. In this work, we developed an *in vitro* osmotic gradient experimental system with increasing osmotic potentials. The system generates a controlled osmotic gradient in the root growth zone while exposing the aerial tissues to control conditions. The osmotic gradient system allowed Arabidopsis seedlings of Col-0 and *ttl1* mutant to sustain proper root growth for 25 days, reaching osmotic potentials of −1.2 MPa. We demonstrated that roots of seedlings grown in the osmotic gradient sustain a higher root growth rate than those that were grown under a homogeneous high osmotic potential. Furthermore, we found out that the expression of some genes is modified in the roots grown in the osmotic gradient compared to those grown in osmotic shock. Our data indicate that using an osmotic gradient can improve our understanding of how plants respond to osmotic stress and help find new genes to improve plant field performance.

## 1. Introduction

Roots directly encounter fluctuating water potential in soil. The soil’s heterogeneous nature, including air pockets and hydrated particles, exposes roots to varying water availability during exploration. Roots detect local water conditions and initiate cellular responses, while long-distance systemic signals coordinate multicellular adjustments, impacting root growth (Gorgues et al., 2022).

Primary root growth in *Arabidopsis thaliana* is governed by a limited number of stem cells within the root apical meristem (RAM), responsible for producing all root cell types through controlled cell division, followed by regulated cellular expansion and differentiation (Scheres et al., 2002, Svolacchia et al., 2020). The Arabidopsis primary root is longitudinally organized into four developmental zones: the proximal meristem (PM), transition zone (TZ), elongation zone (EZ), and differentiation zone (DZ) (Cederholm et al., 2012; Salvi et al., 2020). The PM spans from the quiescent center (QC) to the first elongated cell, where isodiametric cells with high mitotic activity initiate radial expansion, determining root width (Cederholm et al., 2012). In the TZ, cells near the PM maintain mitotic activity, while distal cells undergo anisotropic expansion along the longitudinal axis (Cederholm et al., 2012). Within the EZ, cells elongate exponentially and are characterized by large vacuoles and nuclei displaced toward the cell wall. Various cellular changes occur in this zone, including microtubule reorientation, cell wall softening, and new cell wall synthesis (Petricka et al., 2012). Fully elongated cells transition to the DZ, where they terminally differentiate and cease elongation (Verbelen et al., 2006; Cederholm et al., 2012).

After 7 days from germination, root growth enters a stationary phase, resulting from the delicate balance of cell division, elongation, and differentiation, wherein the regulation of the direction and extent of cell wall expansion in EZ cells plays a pivotal role (Chaiwanon et al., 2016; Pavelescu et al., 2018). During the stationary phase, the sizes of the PM and EZ remain constant, and root growth proceeds by lengthening the DZ, influenced by mature cell length and the rate of cell proliferation in the meristem (Verbelen et al., 2006; Pavelescu et al., 2018).

The meristematic cells are characterized by having thin primary cell walls (0.1-1 μm thick) consisting mostly of complex polysaccharides and a small number of proteins (Cosgrove, 2005; Höfte and Voxeur, 2017). The generic composition of the primary cell wall of Arabidopsis can be described as: 15-40% cellulose, 20-30% hemicellulose, and 30-50% pectins(Cosgrove and Jarvis, 2012). Of the three components, cellulose possesses high tensile strength that prevents and/or directs cell expansion. Cellulose is synthesized by a membranous protein complex part of the CELLULOSE SYNTHASE A (CESA) family, which moves using microtubules as a guide (Cosgrove, 1997). For the synthesis of the primary cell wall, CESA1, CESA3, and CESA6 are required (Cosgrove, 2005). While the presence of the cell wall confers advantages such as a robust exoskeleton and protection against biotic and abiotic factors, it poses a challenge for cell expansión(Gorgues et al., 2022). For plant cells to expand, the structure of the cell wall must loosen (Cosgrove, 2005), a process mediated by proteins called α- EXPANSINS (EXPAs) located in the cell wall itself and activated by a low pH in the apoplast (Cosgrove, 2005, 2015; Sampedro and Cosgrove, 2005). Expansins break hydrogen bonds between cellulose microfibrils and between these and other cell wall components (Cosgrove, 2000; Quiroz-Castañeda & Folch-Mallol, 2011), thus enabling turgor-driven cell expansion. It has been demonstrated that in the Arabidopsis root meristem, proton pumps anchored in the membranes AHA1 and AHA2 create the necessary pH conditions for the protein EXPA1 to loosen the cell walls and, consequently, cell elongation occurs (Pacifici et al., 2018). Once elongated, the cell walls must be resynthesized by incorporating new polymers or modulating their interaction to maintain the shape and size achieved. This mechanism is essential during acclimatization to osmotic or saline stress (Rui and Dinneny, 2019). Any change in osmotic gradient, either by increasing or decreasing the water potential in the external medium or changes in the internal concentration of solutes, leads to direct changes in turgor that can modify cell volume and tissue rigidity. Therefore, to allow growth by expansion, cells must maintain a constant dialogue between the physical properties of the cell wall, solute concentration, and turgor (Gorgues et al., 2022). The exact nature of the signals that allow plants to perceive water availability changes is still unclear. At the cellular level, changes in external osmolarity directly impact the mechanical properties of the cell wall and/or membrane (Gorgues et al., 2022).

Over the past few years, numerous studies have underscored the significance of brassinosteroids (BR) in orchestrating root growth(González-García et al., 2011; Hacham et al., 2011; Geng et al., 2013; Vilarrasa-Blasi et al., 2014; Chaiwanon and Wang, 2015). Mutants lacking BR exhibit shorter roots which is attributed to the diminished elongation of mature cells. Additionally, treatments with elevated BR concentrations impede root growth by diminishing meristem size through accelerated cellular elongation (Chaiwanon and Wang, 2015). The BRs promote mitotic activity and the expression of *CYCLIN D3* and *CYCLIN B1* in the root meristem (González-García et al., 2011; Hacham et al., 2011). Perception of BRs in the epidermis is sufficient to control root growth and meristem size (Hacham et al., 2011). During the cell elongation phase, elevated cellulose synthesis is necessary for cell wall reconstruction(Refrégier et al., 2004). Mutants deficient in BR synthesis or perception have been observed to contain less cellulose than wild-type genotypes(Xie et al., 2011). The expression of *CESA* genes related to primary cell wall synthesis is decreased in these mutants and is only induced by external BR application in mutants deficient in hormone synthesis (Xie et al., 2011). Furthermore, it is known that the transcription factor BES1 can associate with promoters of most *CESA* genes related to primary wall synthesis (Xie et al., 2011). During primary root growth, BR signaling activates BES1 transcription and promotes the association of BES1 with E-box motifs (CANNTG) in the promoters of *CESA1*, *CESA3*, and *CESA6* genes to activate their expression and cellulose synthesis in elongation zone cells, thereby promoting growth (Xie et al., 2011; Novaković et al., 2018).

*TETRATRICOPEPTIDE THIOREDOXIN-LIKE (TTL)* genes encode a unique family of proteins found specifically in land plants, characterized by six tetratricopeptide repeat (TPR) domains situated at specific positions within the sequence, as well as a thioredoxin-homologous sequence in the C-terminal position (Rosado et al., 2006; Lakhssassi et al., 2012). TPR domains are well-known protein-protein interaction modules. Mutations in the Arabidopsis *TETRATRICOPEPTIDE THIOREDOXIN-LIKE 1* (*TTL1*) gene results in root swelling and growth arrest under NaCl and osmotic stress (Rosado et al., 2006; Lakhssassi et al., 2012; Amorim-Silva et al., 2019). We previously demonstrated by atomic force microscopy that the *ttl1* mutant has more elastic cell walls than the wild-type genotype in epidermal cells of root EZ, which explains the swelling phenotype observed in the mutant (Cuadrado-Pedetti et al., 2021). This evidence, together with the confirmation of genetic interaction between *TTL1* and *CESA6*, a component of the *CELLULOSE SYNTHASE COMPLEX* (*CSC*) responsible for synthesizing primary cell walls in the root apical meristem, highlighted the role of *TTL1* in maintaining cell wall integrity (Amorim-Silva et al., 2019; Cuadrado-Pedetti et al., 2021). (Amorim-Silva et al., 2019) have proved in vivo interaction between TTL3 and constitutively active BRASSINOSTEROID INSENSITIVE 1 (BRI1), BRI SUPPRESOR 1 (BSU1), and BRASSINOZOLE-RESISTAMT 1 (BZR1) and that TTL3 exhibits dual cytoplasmic and membrane localization, which is dependent on endogenous brassinosteroid (BR) content, suggesting that TTL proteins may act as positive regulators of BR signaling (Amorim-Silva et al., 2019). Furthermore, the suppression of *TTL1* expression and the BR signaling pathway in response to osmotic stress and BR treatment (Amorim-Silva et al., 2019) and the interaction of TTL1 with CESA1 (Kesten et al., 2022) suggests a role for TTL1 in mediating adaptation to osmotic stress through auxin and brassinosteroid homeostasis (Cuadrado-Pedetti et al., 2021).

In this study, we undertook a novel approach to evaluate Arabidopsis primary root growth. We simulated a scenario where the root tip gradually encounters increasing osmotic potentials, creating an osmotic gradient condition. This system, unique in its design, generates a controlled osmotic gradient in the root growth zone while maintaining the aerial tissues under control conditions. Our findings revealed that this osmotic gradient system enabled Arabidopsis seedlings of Col-0 and *ttl1* mutant to sustain proper root growth for 25 days, reaching osmotic potentials of −1.2 MPa. Notably, roots of both genotypes grown in the osmotic gradient exhibited a higher root growth rate than those grown under a homogeneous high osmotic potential. Intriguingly, when *ttl1* seedlings were grown in the osmotic gradient, they did not exhibit the characteristic swelling phenotype in the root at the extreme osmotic potential (−1.2 MPa).

Furthermore, our study revealed that the osmotic gradient had a significant impact on the expression of genes of the *TTL* family expressed in the primary root (*TTL1* and *TTL3*); primary cell wall cellulose synthesis-related genes: *CESA1*, *CESA3*, *CESA6*; a mannan synthase: *CESA-LIKE 9 (CSLA9)*(Zhu et al., 2003; Davis et al., 2010); a gene related to cellulose microfibrils deposition and crystallization: *COBRA;* genes related to cell wall remodeling during anisotropic expansion: *EXPANSIN A1*, *AHA1*, and *AHA2*, genes related to brassinosteroid synthesis and signaling: *DWARF4* (*DWF4*), *CONSTITUTIVE PHOTOMORPHOGENIC DWARF* (*CPD*), and genes related to the cell cycle and cell expansion: *CYCLIN D3;1* (*CYCD3;1*). Notably, 10 of the 14 genes assayed by RT-qPCR showed significant genotype by osmotic treatment interaction. Among these genes, we observed three distinct patterns of expression in Col-0 and *ttl1.* One pattern was the induction of the expression levels in both genotypes grown in the osmotic treatments compared to control conditions, reaching a higher expression level in the osmotic gradient than in the osmotic shock. The second pattern was of genes that showed repression in Col-0 grown in both osmotic treatments compared to the expression in Col- 0 grown in control conditions and induction in *ttl1* grown in osmotic shock compared to the expression in *ttl1* grown in control and osmotic gradient. A third group of genes presented a pattern of expression induction in Col-0 grown in the osmotic gradient and a repression in the osmotic shock compared to the expression observed in Col-0 grown in control conditions. However, this group of genes showed an expression induction in *ttl1* grown in the osmotic gradient and the osmotic shock compared to the expression in *ttl1* grown in control conditions. These results suggest that the osmotic gradient could significantly enhance our understanding of how roots respond to osmotic stress and could potentially improve the translation of data generated in vitro to field trials, offering a promising avenue for future research.

## 2. Results

In this study, we developed an osmotic gradient experimental system (Supplemental Figure 1) with increasing osmotic potentials generated with solid MS + 1.5% sucrose containing from 0 to 400 mM mannitol (corresponding to −0.4 to −1.2 Mega Pascal (MPa) of osmotic potential) to investigate root adaptation responses to local water availability.

### Evaluation of the Osmotic Gradient System

To address the effectiveness of the osmotic gradient system, we conducted a seed germination assay using *ttl1* (a mutant known for increased germination under osmotic stress conditions; Rosado et al., 2006) and its background line Col-0 (Supplemental Figure 2). The seeds were placed in each mannitol concentration strip of the osmotic gradient (Supplemental Figure 2 B) and the percentage of germinated seeds was calculated relative to the total number of seeds plated in each mannitol concentration strip (Supplemental Figure 2C). We observed a negative correlation between the increasing osmotic potential and the percentage of seed germination for both genotypes (correlation for Col-0 r= −0,92 p= 5,75281E-33; correlation for *ttl1* r= −0,96 p= 2,69247E-45). Interestingly, at 400 mM of mannitol, Col- 0 exhibited an 80% reduction in germination while *ttl1* showed a 60% reduction (Supplemental Figure 2B and C). These finding agrees with the increased germination response in osmotic stress reported for *ttl1* (Rosado et al., 2006). In summary, our results demonstrate that the osmotic gradient system, with increasing osmotic potential, is suitable for primary root growth studies.

### The reduction in root growth rate of Col-0 and *ttl1* grown in the osmotic gradient was milder than in the osmotic shock conditions

The root length relies on the equilibrium between the rate of cell division and cell elongation (Cederholm et al., 2012; Chaiwanon and Wang, 2015). Typically, a more significant number of cells dividing at the root apical meristem (RAM) leads to more cells available for elongation and differentiation, thus contributing to a higher rate of root growth (Baskin, 2013; Salvi et al., 2020; Svolacchia et al., 2020).

This work aims to evaluate the primary root growth under osmotic gradient conditions. For this purpose, we grew Col-0 and *ttl1* seven days post-germination seedlings in solid MS + 1.5% sucrose containing a range from 0 to 400 mM mannitol corresponding to −0.4 to −1.2 MegaPascal (MPa) of osmotic potential (Supplemental Figure 1) and compared root growth behavior in the osmotic gradient for both genotypes with the ones obtained by(Cuadrado-Pedetti et al., 2021) under osmotic shock conditions (Figure 1). Interestingly, root growth rates of seedlings grown in the osmotic gradient slowed down less than in osmotic shock. After 25 days of growth in the osmotic gradient, when the roots reached the range of −1 to −1.2 MPa of osmotic potential, the root growth rate of Col-0 (2.2 ± 0.2 mm/day; Figure 1) was 44% lower than the one observed under control conditions (Col-0: 3.95 ± 0.1 mm/day; Figure 1). However, the *ttl1* mutant experienced a reduction of 62% of its root growth rate (1.5 ± 0,2 mm/day; Figure 1) after growing for 25 days in the osmotic gradient compared to Col-0 grown in control conditions (Figure 1). In contrast, the reduction in root growth rate observed after 7 days in an osmotic shock medium at −1.2 MPa was 88% for Col-0 and 95% for *ttl1,* compared to the root growth rate observed under control growth conditions (Cuadrado-Pedetti et al., 2021).

**Figure 1.**
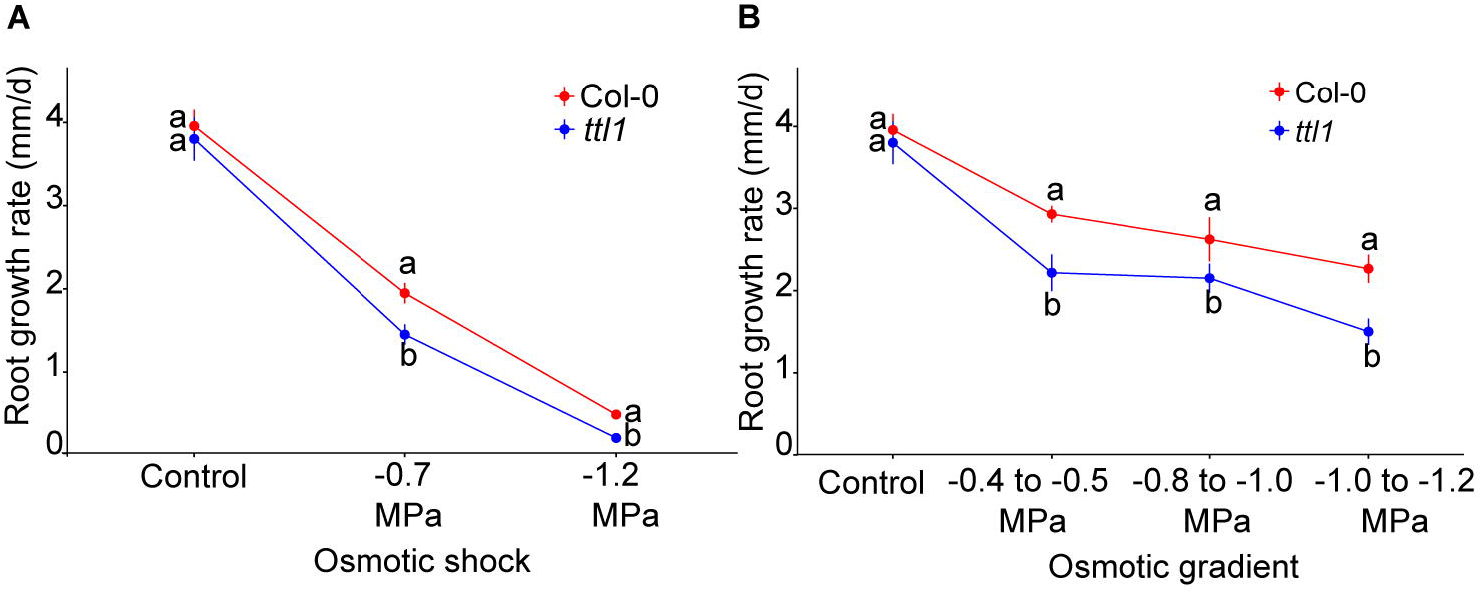
The osmotic gradient system allowed Arabidopsis seedlings of Col-0 and *ttl1* mutant to sustain proper root growth for 25 days. (**A**) The graphic shows the root growth rate for Col-0 and *ttl1* grown in osmotic shock conditions (−1.2 MPa). (**B**)The graphic shows the root growth rate for Col-0 and *ttl1* grown during 25 days in an osmotic gradient with an increasing osmotic potential from −0.4 MPa to −1.2MPa. Although the increasing osmotic potentials decelerate root growth rate, both genotypes maintain a greater root growth rate at −1.2MPa in comparison to the osmotic shock condition. The magnitude of deceleration is higher in *ttl1*. Different letters indicate statistically significant differences (t Student *p* < 0.001; n = 30).

We determined the growth parameters influencing the rate of root growth, specifically in the cortical cells extending from the stem cell initials near the quiescent center to the onset of the differentiation zone in roots grown during 25 days in the osmotic gradient (Perilli and Sabatini, 2010; Cole et al., 2014). The characterization in control conditions showed that the *ttl1* mutant has a reduced number of cortical cells in the PM compared to Col-0; moreover, *ttl1* cells at maturity did not elongate to the same extent as Col-0 cells (Figure 2A) as described before by (Cuadrado-Pedetti et al., 2021). After 25 days of growth in the osmotic gradient, both genotypes significantly reduced the number of cells in the PM, Col-0 by 49% and *ttl1* by 51% (Figure 2A) compared to the cell number in control conditions. However, the reduction was milder compared to what was observed after 7 days of growth at an osmotic shock medium with an osmotic potential of −1.2 MPa, where a Col-0 experienced a 72% and *ttl1* a 76% reduction in the cell number (Cuadrado-Pedetti et al., 2021). Moreover, the mature cell length of roots of both genotypes grown in osmotic gradient conditions experienced a less pronounced reduction in length than what was observed in the osmotic shock medium (Figure 2B).

**Figure 2.**
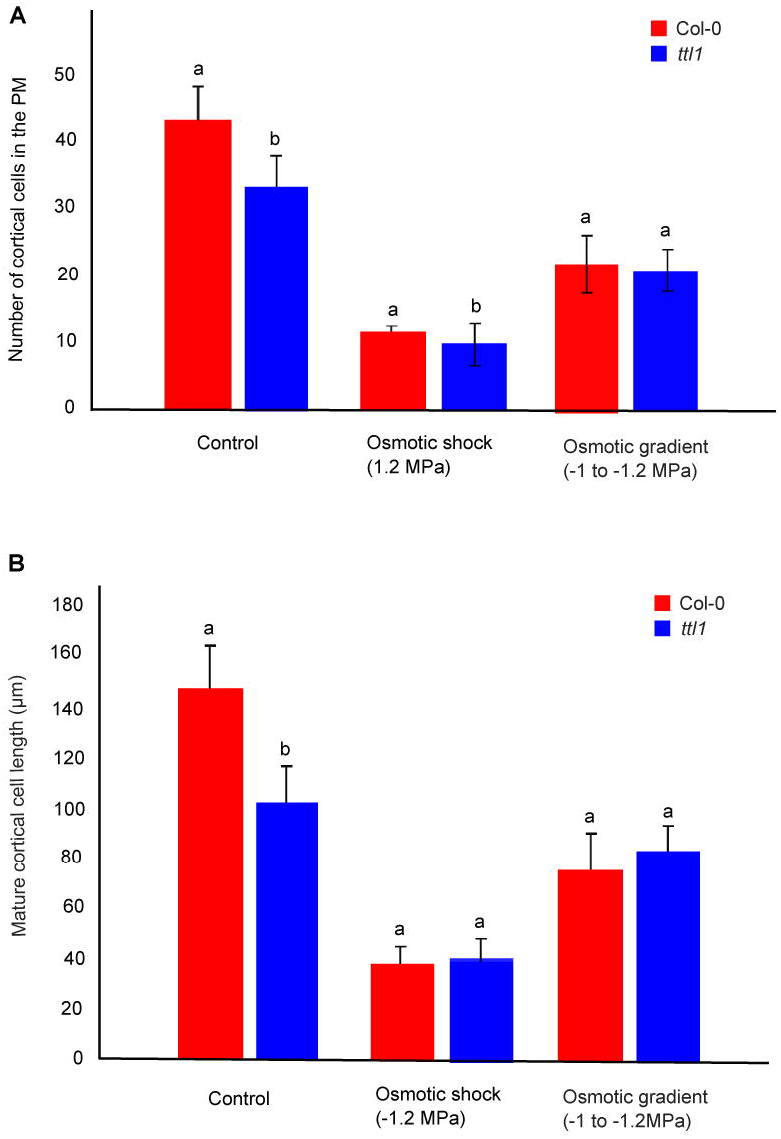
The figure shows the cortical cell number in the proximal meristem (A) and the length of mature cortical cells (B) for both Col-0 and *ttl1* grown in control and in an osmotic gradient with an increasing osmotic potential from −0.4 MPa to −1.2Mpa. Data are presented as means ± standard error (SE). Different letters denote statistically significant differences (Student’s t-test, p < 0.05; n = 10). The experiments were repeated three times.

The cortical cell profile analysis described in Materials and Methods was used to estimate the cell production rate, the cell cycle lenght and the average time between each cell entering the TZ and the EZ.

In control conditions, the cell production rate is higher in *ttl1*, the length of the cell cycle is shorter and the average time between each cell entering the TZ and the EZ was faster compared to Col-0, as already reported by Cuadrado-Pedetii et al. 2021. When the parameters were analyzed for roots of *ttl1* grown in the osmotic gradient we observed an inversion in the pattern of these parameters compared to control conditions (Figure 3) and to what was reported in osmotic shock by Cuadrado-Pedetti et al. 2021. The osmotic gradient led to a decrease in the cell production rate (Figure 3A), an increase in the length of the cell cycle (Figure 3 B) and an increase in the average time between each cell entering the TZ and the EZ in roots of *ttl1* compared to Col-0 (Figure 3C). Interestingly, we did not see the swelling phenotype in *ttl1* roots growing in the osmotic gradient at the extreme osmotic potential (−1.2 MPa).

**Figure 3.**
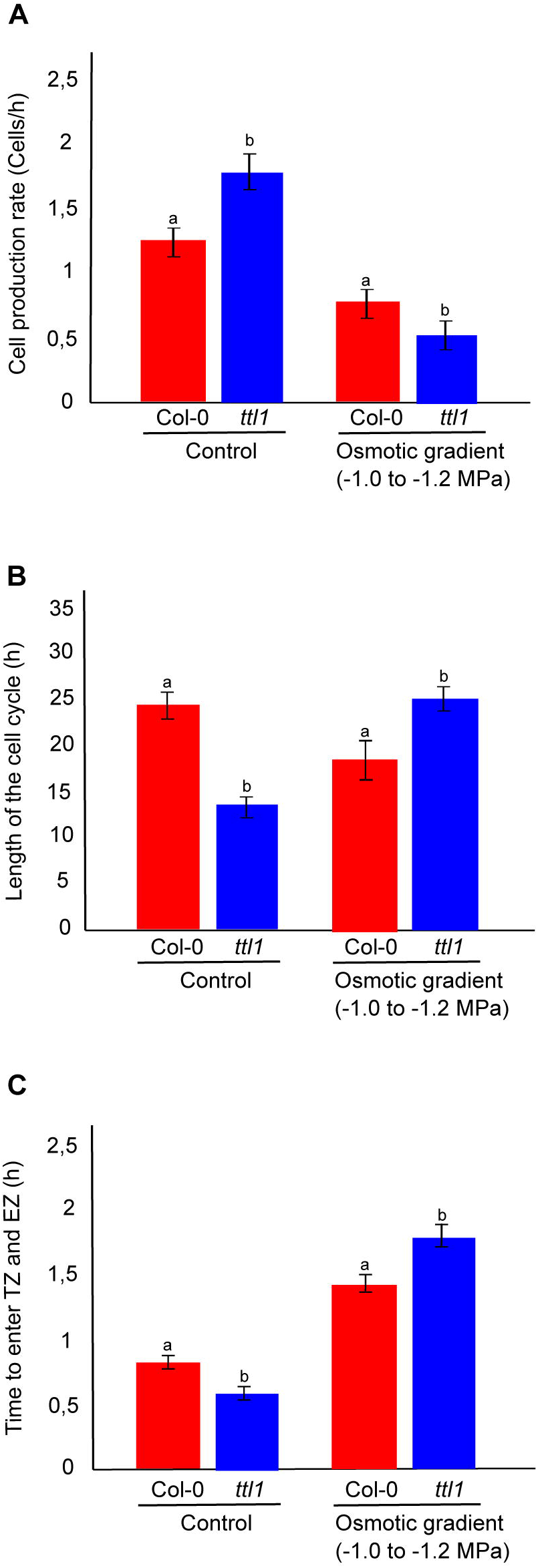
The figure shows the growth parameters in the PM of Col-0 and *ttl1* roots grown in control and osmotic gradient conditions. The figure shows the cell production rate (cells/h) (**A**); the length of the cell cycle (h) (**B**); and the time to enter the transition zone (TZ) and elongation zone (EZ) (**C**) for Col-0 and *ttl1* grown during 7 days in control and in an osmotic gradient from −0.4 to −1.2 MP. Values are means ± standard error (SE). Different letters indicate statistically significant differences (*t* Student *p* < 0.05; *n* = 10).

The expression of genes involved in anisotropic cell expansion were significantly downregulated in roots of the *ttl1* mutant and presented a distinct pattern of modulation depending on the stress experimental condition.

To delve deeper into understanding the Col-0 and *ttl1* root responses to the different experimental conditions (control, osmotic shock and osmotic gradient), we used quantitative RT-PCR (qRT-PCR) to analyze changes in gene expression of key genes orchestrating root growth: genes of the *TTL* family expressed in the primary root (*TTL1* and *TTL3*); primary cell wall Cellulose synthesis-related genes: *CESA1*, *CESA3*, *CESA6*; a mannan synthase: *CSLA9*; genes related to cell wall remodeling during anisotropic expansion: *PECTATE LYASE 12* (*PLL12*), *Expansin A1*, *AHA1*, and *AHA2*; genes related to brassinosteroid synthesis and signaling: *DWARF4* (*DWF4*), *CONSTITUTIVE PHOTOMORPHOGENIC DWARF* (*CPD*), *BRASSINAZOLE-RESISTANT 1* (*BES1*); genes related to the cell cycle and cell expansion: *CYCLIN D3;1* (*CYCD3;1*).

Exploration of the RT-qPCR data using distance matrix analysis (heatmap) showed that most of the biological replicates of both genotypes clustered within each experimental condition group, despite one replica of *ttl1* grown in osmotic gradient that clustered outside its experimental condition group. Interestingly, the replicas of the osmotic shock and control experimental conditions clustered together, whereas the osmotic gradient replicas formed a separate cluster (Figure 4 A). This was also observed when we performed a principal component analysis (PCA), from which we obtained six well-defined groups corresponding to each genotype grown in the three osmotic conditions (control: green ellipses; osmotic gradient: pink ellipses; and osmotic shock: yellow ellipses (Figure 4 B). The principal component 1 (PC1) explained the greatest proportion of the variance (46%) separating the samples grown in control condition (green ellipse) and the samples grown in osmotic gradient (pink ellipse). The samples of Col-0 grown in osmotic shock (yellow ellipse, circle dots) did not differ from the ones grown in control condition regarding PC1 (Figure 4 B). The genes that correlated the most with PC1 were *AHA1*, *AHA2*, *CESA1*, *CESA3*, *CESA6*, *CYCD3;1*, *PLL12*, and *TTL3* with correlations values over 0.70 (Supplemental Table 1), meaning that the expression of these genes was greater in the osmotic gradient replicas. The principal component 2 (PC2), which explained 24.4% of the variance, clearly separated the samples grown in control conditions (green ellipses) by genotype (circles (Col-0) vs. triangles (*ttl1*)). *CSLA9*, *COBRA*, *CPD*, and *DWF4* were the most correlated genes with PC2 with correlation values lower than −0.70 (Supplemental Table 3), meaning that the expression levels of these genes are greater in samples of Col-0 grown in control conditions compared to samples of *ttl1* grown in the same condition and the samples of both genotypes grown in other experimental conditions. (Figure 4 B).

**Figure 4.**
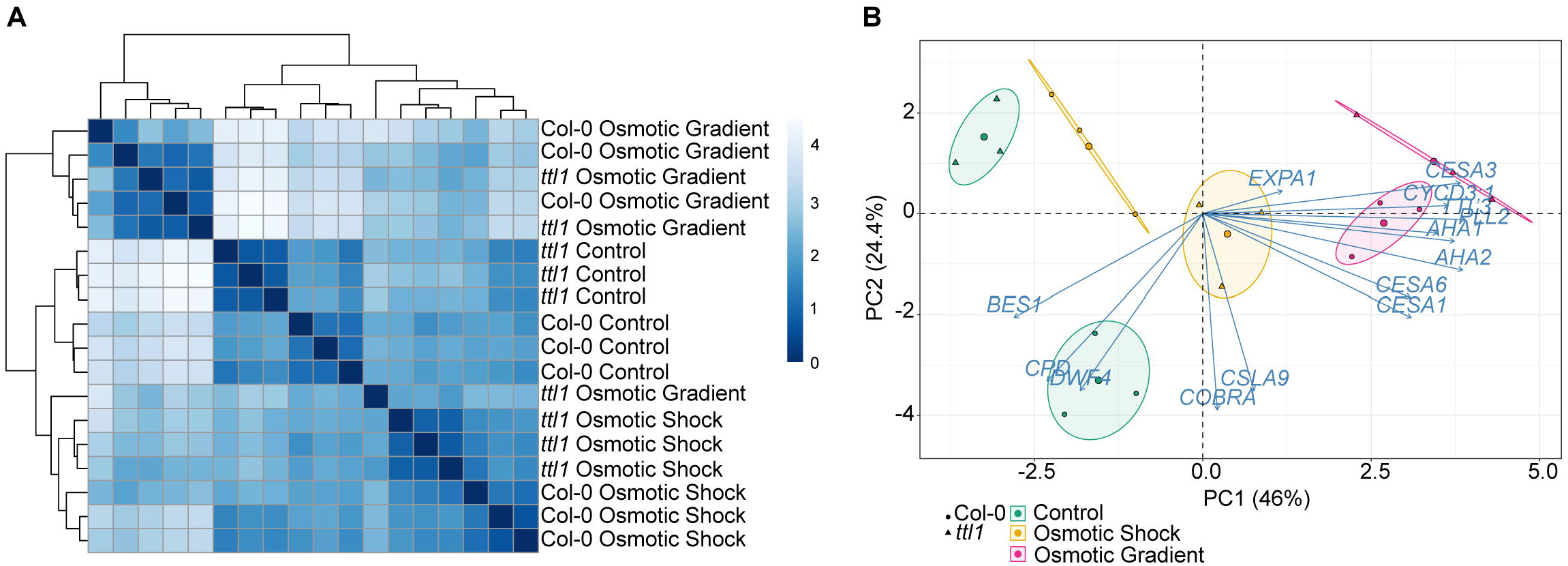
Descriptive analysis of the RT-qPCR data **(A)** Heatmap of all samples comprising three replicates of each genotype (Col-0 and *ttl1*) and growing conditions (Control, osmotic shock, osmotic gradient). (**B**) Principal component analysis (PCA) of gene expression levels in Arabidopsis thaliana roots that had been grown in control, osmotic shock, and osmotic gradient conditions. Colors indicate the growing condition: Green for control, orange for osmotic shock, and pink for the osmotic gradient. The circle indicates Col-0 and the triangle *ttl1* mutant. Percentages of variation explained by each PC are indicated along the axes.

The clustering analysis made using the average expression of each gene in each genotype and experimental condition showed three clusters (Figure 5) with genes with a distinct expression pattern in Col-0 and *ttl1.* The first cluster contains only *CYCD3;1* that showed induction of the expression levels in both genotypes grown in the osmotic treatments compared to what was observed in both genotypes grown in control conditions, the induction being greater in the osmotic gradient than the osmotic shock (Figure 5). The second cluster contains five genes (*DWF4*, *BES1*, *CPD*, *CSLA9,* and *COBRA*) which its expression was repressed in Col-0 grown in both the osmotic gradient and the osmotic shock compared to the expression in Col-0 grown in control conditions and a pattern of induction in *ttl1* grown in osmotic shock compared to the pattern of expression in *ttl1* grown in control and in the osmotic gradient conditions (Figure 5); except for *CPD* and *DWRF4* which expression was repressed in ttl1 roots grown in osmotic shock conditions. The third cluster contains 8 genes (*AHA2*, *AHA1*, *CESA1*, *CESA3*, *CESA6*, *EXPA1*, *PLL12,* and *TTL3*) that are characterized by a pattern of expression induction in Col-0 grown in the osmotic gradient compared to the expression observed in Col-0 grown in control and osmotic shock conditions. Moreover, the third cluster showed a pattern of expression induction in *ttl1* grown in the osmotic gradient and the osmotic shock compared to the pattern of expression in *ttl1* grown in control conditions.

**Figure 5.**
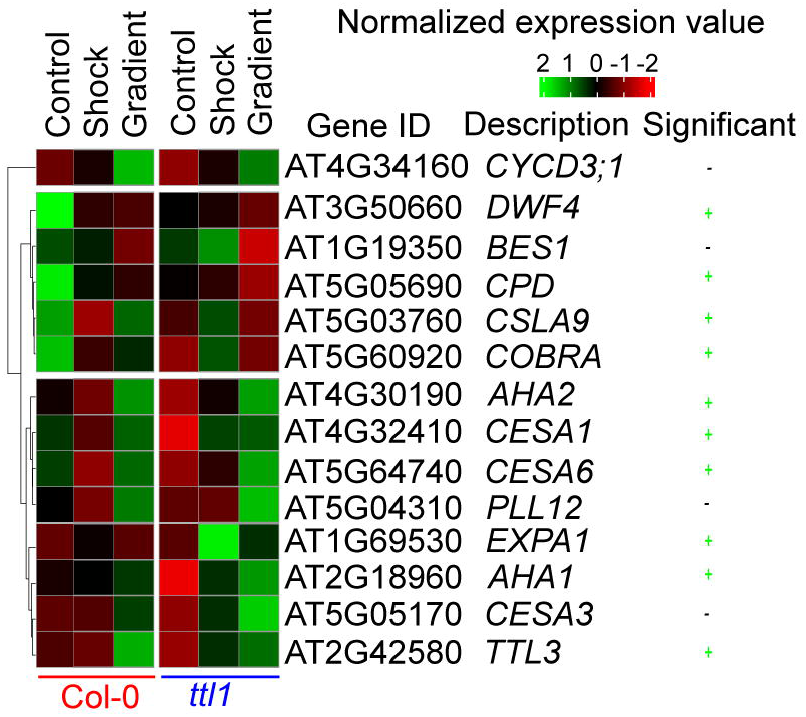
A heat map showing the normalized expression of genes in the Col-0 or *ttl1* genetic backgrounds after transfer to control, osmotic shock and osmotic gradient conditions. Expression measured using RT-qPCR from RNA isolated from whole roots. A two-way ANOVA was used to identify genes showing significant genotypes by treatment effects (marked in green, P value < 0.05).

Of the 14 genes assayed, we found that ten genes showed significant genotype by osmotic treatment effects (Figure 5; two-way analysis of variance (ANOVA), multiple-testing corrected P value < 0.05; Figure 5).

Interestingly, we found that in control conditions, *ttl1* mutant showed statistically significant downregulated expression of the following genes *CYCD3;1*, *CPD, DWF4, CESA1, CESA6, COBRA, AHA1, and AHA2* compared to what was observed in roots of Col-0 grown in this condition (Figure 6; single way analysis of variance (ANOVA), P value < 0.05; Supplemental Figure 3). Both osmotic stress conditions downregulated the expression of *CPD* and *DWF4* in roots of Col-0 to the levels found in *ttl1* mutant growing in control conditions (Figure 6 A and B). However, we did not observe changes in the levels of the expression of these genes in roots of the *ttl1* mutant due to the osmotic stress condition assayed (Figure 6 A and B). The expression of *CESA1* did not change in Col-0 roots grown in osmotic shock conditions compared with the expression levels observed in roots of Col-0 grown in control conditions. However, the expression of *CESA1* was upregulated in roots of Col-0 grown in the osmotic gradient (Figure 6C). Notable, the expression levels of *CESA1* in roots of the *ttl1* mutant were upregulated by the osmotic shock to the levels of expression observed in roots of Col-0 grown in control conditions and reached higher values of expression in roots of *ttl1* grown in the osmotic gradient (Figure 6 C). The osmotic shock condition had a different effect on the expression levels of *CESA6* than the osmotic gradient in roots of Col-0. The osmotic shock downregulated *CESA6* in roots of Col- 0 compared to the expression levels observed in Col-0 grown in control conditions (Figure 6 D). However, in the osmotic gradient condition *CESA6* expression levels stayed as the levels observed in Col-0 in control conditions (Figure 6 D). On the contrary, both osmotic shock and osmotic gradient upregulated *CESA6* in the roots of *ttl1,* although *CESA6* reached the highest expression levels in the osmotic gradient (Figure 6 D). Also, we observed a differential effect of the osmotic condition in the expression levels of *CSLA9* depending on the genotype evaluated. While in the roots of Col-0 *CSLA9* was downregulated by both the osmotic shock and the osmotic gradient, in the roots of *ttl1*, its expression levels stayed unchanged independent of the growth condition (Figure 6 E). *COBRA* expression was downregulated by the osmotic gradient and the osmotic shock conditions in roots of Col-0 (Figure 6 F). However, its expression was upregulated by the osmotic shock in roots of *ttl1* and maintained constant in response to the osmotic gradient, compared to its expression in control conditions (Figure 6 F). We did not find statistically differences between the expression levels of *TTL1* in Col-0 grown in the different growing conditions (Supplemental Figure 3). Both osmotic conditions (shock and gradient) upregulated *TTL3* expression in roots of *ttl1* compared to control conditions. Whilst, in Col-0 roots, the upregulation was observed only in the osmotic gradient condition (Figure 6 G). The expression of *EXPA1* was only upregulated in roots of *ttl1* grown in osmotic shock (Figure 6 H). The expression of *AHA1* was upregulated in roots of Col-0 and *ttl1* in response to both the osmotic shock and the osmotic gradient conditions, reaching higher levels in the second condition (Figure 6 I). Furthermore, the expression of *AHA2* was downregulated in roots of Col-0 grown in osmotic shock and upregulated in roots grown in the osmotic gradient conditions. While in *ttl1* roots, both types of osmotic conditions upregulated the expression of *AHA2* (Figure 6 J).

**Figure 6.**
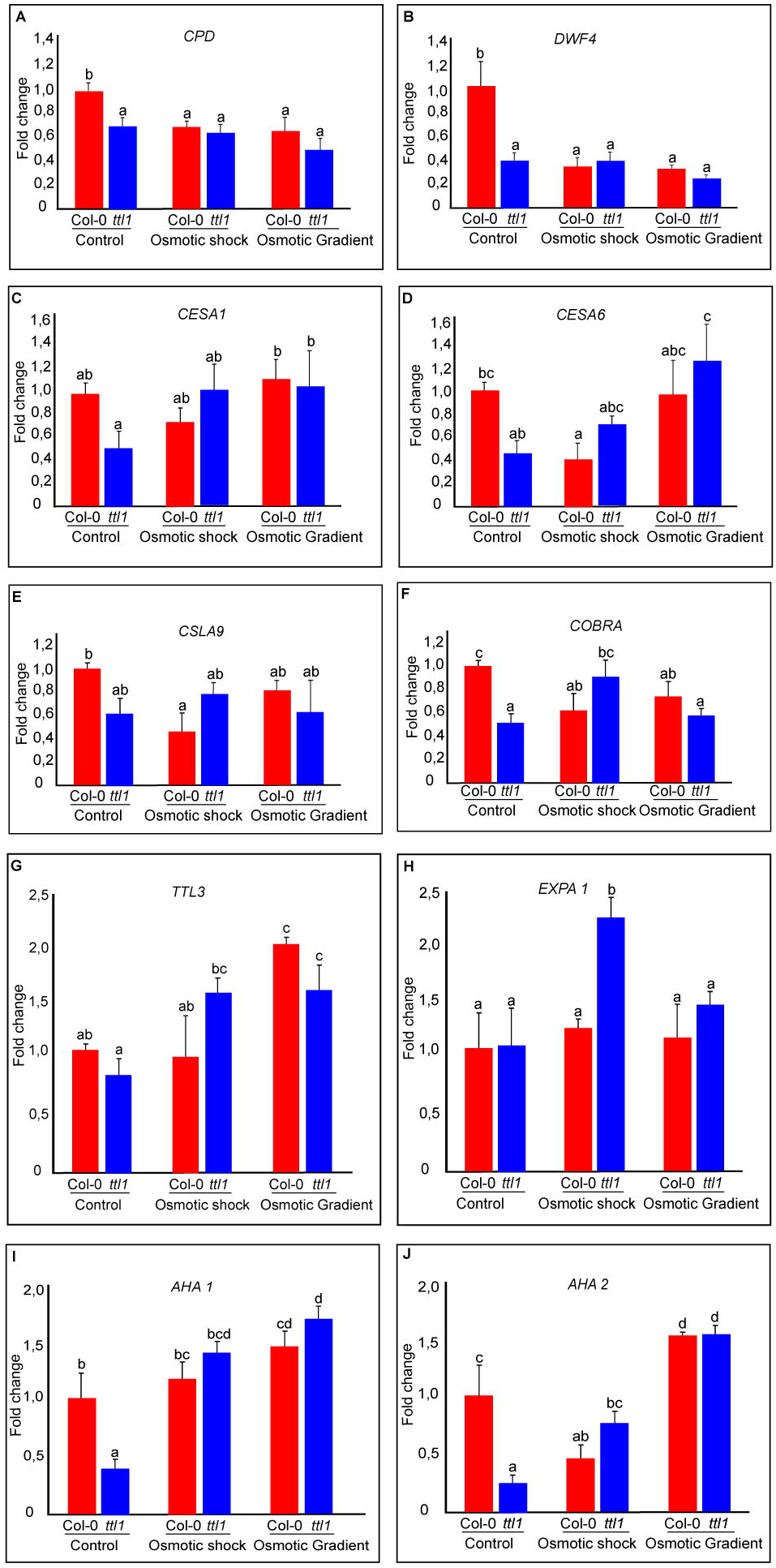
Gene expression of the genes that have significant genotype by environment effects. Data are presented as means ± standard deviation (SD). Three biological replicates and two technical replicates were considered, and gene expression is relative to *CYTOCHROME C OXIDASE RELATED*. Different letters indicate statistically significant differences. A two-way ANOVA followed by the Tukey test was used to identify significant differences (P value < 0.05).

## 3. Discussion

Water is an essential resource that can limit plant growth. Its availability is initially sensed by the root tip tissues (Chang et al., 2024). The ability of roots to adjust their physiology and morphology under water deficit conditions makes this organ a useful model for understanding how plants respond to water stress (Dinneny, 2019; Yuan et al., 2021). In general, osmotic stress induced in vitro by a medium with high homogeneous concentrations of mannitol or polyethylene glycol (osmotic shock) that produced high osmotic potentials reduces primary root growth rate and produces root swelling in hypersensitive mutants (Rowe et al., 2016; Cajero-Sanchez et al., 2019; Cuadrado-Pedetti et al., 2021; Yuan et al., 2021). However, when a root explores the soil, it may not be immersed in a situation where the entire root is exposed to the same osmotic potential. Therefore, in this study, we evaluated Arabidopsis primary root growth using a novel approach where we simulated a situation where the root tip gradually encounters increasing osmotic potentials (osmotic gradient condition). Additionally, a transcriptional analysis of *ttl1*, a mutant related to cell wall integrity maintenance of the primary root meristem, was included in the study.

Plant root growth and development rely on the balance between the rate of cell division, cell elongation, and differentiation. Generally, at the stationary phase, a high number of cells in the meristematic zone (PM) produces more cells that elongate and differentiate, resulting in a higher root growth rate(Salvi et al., 2020; Svolacchia et al., 2020). Using the osmotic gradient condition, we prove that gradually exposing the root tip to increasing osmotic potentials allows the primary root to adapt better to higher osmotic potentials. The osmotic gradient of −0.4 to −1.2 MPa of osmotic potential reduces the root growth rate less than the exposure to an osmotic shock of −1.2 MPa for 7 days. The analysis of the cortical cell profile in the osmotic gradient showed that the PM remains with a higher cell number after the root tip reaches the higher osmotic potential of the gradient compared to what was observed in the osmotic shock condition. Also, we found a genotype-distinct response regarding the different growth parameters analyzed, as was observed before with the osmotic shock condition (Cajero-Sanchez et al., 2019; Cuadrado-Pedetti et al., 2021). Specifically, in roots of *ttl1* grown in control conditions has a higher the cell production rate, the length of the cell cycle is shorter and the average time between each cell entering the TZ and the EZ was faster compared to Col-0 roots, as already reported by (Cuadrado-Pedetti et al., 2021). When the parameters were analyzed for roots of *ttl1* grown in the osmotic gradient we observed an inversion in the pattern of them compared to control conditions (Figure 3) and to was reported in osmotic shock by(Cuadrado-Pedetti et al., 2021). The osmotic gradient led to a decrease in the cell production rate (Figure 3A), an increase in the length of the cell cycle (Figure 3 B) and an increase in the average time between each cell entering the TZ and the EZ in roots of *ttl1* compared to Col-0 (Figure 3C). In the reviewed literature, investigations employing a gradient featuring escalating osmotic potentials generated in the same vial using a gradient maker akin to those delineated in this study are notably absent. Nonetheless, extant studies have documented the utilization of a water gradient characterized by diminishing osmotic potentials (−0.35 MPa to −0.08 MPa, or the exposure of seedlings to homogeneous increasing concentrations of osmotic potentials in different Petri dishes (Saucedo et al., 2012; Miao et al., 2021). Similar results were found comparing homogeneous high temperatures in the root zone of the wild-type Arabidopsis genotype to a gradient of decreasing temperature in the same container. Homogenous high temperatures produced significantly shorter roots with a lower cell division rate in the meristem; although the exposure to the temperature gradient recovered the growth parameters almost to the values observed in standard temperatures (González-García et al., 2023).

Cytokinesis and cell expansion in plants are processes highly dependent on cell wall synthesis and deposition (Engelsdorf et al., 2018; Gigli-Bisceglia and Hamann, 2018). Brassinosteroids and transcriptional regulation have been implicated in cellulose production during primary cell wall formation and cell cycle progression(Gigli-Bisceglia and Hamann, 2018). In Arabidopsis, manipulation of cellulose through inhibition of synthesis or mutations in *CESA* genes activates a set of cell wall damage response genes. These responses include swelling of epidermal cells, changes in cell wall composition and growth inhibition, among others. Mutants with defects in cell wall integrity exhibit enhanced sensitivity to moisture gradients and reduced osmotic tolerance and present significant downregulation of the expression of genes known to encode enzymes catalyzing the biosynthesis of cell wall in the root tips (Chang et al., 2024).

Here, we investigated by RT-qPCR the expression pattern of genes related to cell cycle progression and cell wall deposition during root growth and found out that the gene expression pattern of roots exposed to high osmotic potential is modified when the root systems are grown in an osmotic gradient. Notably, 10 of the 14 genes assayed by RT-qPCR showed significant genotype by environmental effects. Moreover, among these genes, we found distinct patterns of expression in Col-0 and *ttl1.* Interestingly, we found that the *ttl1* mutant has reduced expression levels *CYCD3;1, CPD, DWF4, CESA1, CESA6, CSLA9, COBRA, TTL3, AHA1, and AHA2* in roots grown in control conditions in agreement with what was expected considering the previously reported *ttl1* phenotypes (more elastic cell walls and fewer cells in the PM) in control conditions. The osmotic treatment induced the expression level of *CYCD3;1* in both genotypes compared to what was observed in control conditions, the induction being greater in the osmotic gradient than in the osmotic shock (Figure 5). *DWF4*, *BES1*, *CPD*, *CSLA9*, and *COBRA* showed a repression pattern in Col-0 grown in both the osmotic gradient and the osmotic shock compared to the expression in Col-0 grown in control conditions. However, these genes showed a pattern of induction in *ttl1* grown in osmotic shock compared to the pattern of expression in control and osmotic gradient conditions (Figure 6). A third group of genes, *AHA2*, *AHA1*, *CESA1*, *CESA3*, *CESA6*, *EXPA1*, *PLL12*, and *TTL3*, were characterized by a pattern of expression induction in Col-0 grown in the osmotic gradient and a pattern of repression in the osmotic shock compared to the expression pattern observed in Col-0 grown in control conditions. Moreover, this group of genes showed a pattern of expression induction in *ttl1* grown in the osmotic gradient and the osmotic shock compared to the pattern of expression in *ttl1* grown in control conditions (Figure 5 and 6). The differential clustering of genes in distinct patterns of expression put on evidence that homogeneous osmotic stress compromises the modulation of gene expression and root growth adaptability as was also reported by (González-García et al., 2023) with homogeneous temperatures.

Our data indicate that using an osmotic gradient can improve our understanding of how plants respond to osmotic stress and help find new genes to improve plant field performance.

## Conclusion

Proper growth of the primary root involves auxin/BR homeostasis and cell wall integrity (Chaiwanon and Wang, 2015). During the cell elongation phase, elevated cellulose synthesis is necessary for cell wall reconstruction (Refrégier et al., 2004). The expression of *CESA* genes is downregulated in mutants impaired in BR synthesis (Xie et al., 2011). Furthermore, it is known that the transcription factor BES1 can associate with promoters of most *CESA* genes (Xie et al., 2011) to activate their expression and cellulose synthesis in elongation zone cells, thereby promoting growth (Xie et al., 2011; Novaković et al., 2018). The *ttl1* mutant has been described as a mutant affected in auxin/BR homeostasis that has a more elastic cell wall in cells of the EZ in roots grown in control conditions (Cuadrado-Pedetti et al., 2021). Our RT-qPCR data showed that in roots of *ttl1* grown in control conditions, the expression of *CYCD3;1*, *CPD, DWF4, CESA1, CESA6, COBRA, AHA1, and AHA2* was downregulated compared to what was observed in roots of Col-0, which is consistent with the previously reported phenotypes regarding cell wall integrity and cellular number in the PM of the mutant. The downregulation of the expression of *CPD* and *DWF4* suggests lower contents and the unchanged expression levels of *BES1* compared to Col-0 suggesting an active BR signaling pathway in roots of the *ttl1* mutant grown in control conditions. The differential expression of these genes in the *ttl1* mutant compared to Col-0, together with the cytoplasm localization of TTL1, makes us hypothesize that in the *ttl1* mutant a transcript differential degradation rate may be occurring. This possible regulatory mechanism may involve the TTL1 by complexing with other proteins, stabilizing the transcripts of these genes and avoiding its degradation (Figure 7).

**Figure 7.**
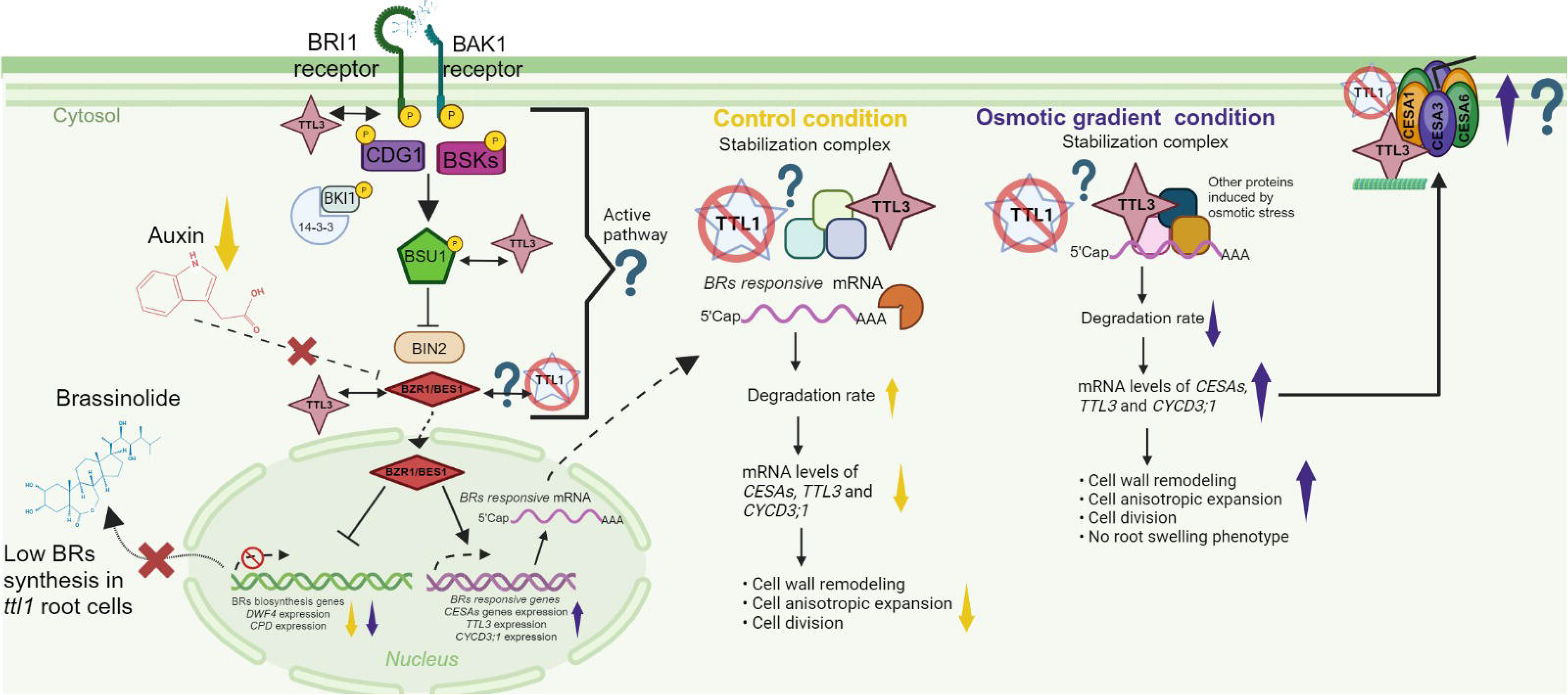
A model that illustrates a transcriptional regulatory scenario for *ttl1* primary root growth in control and osmotic gradient conditions. Primary root growth depends on the Auxin/Brassinosteroid homeostasis and proper cell wall remodeling. The model hypothesizes that the TTL1 protein could play a role in stabilizing specific mRNAs in the cytoplasm preventing its degradation. The yellow arrows indicate the levels of transcripts of the genes or processes specified in the control condition. The violet arrows indicate the levels of transcripts of the genes or processes specified in the osmotic gradient condition. The abbreviations indicate these genes or proteins: BRASSINOSTEROID INSENSITIVE 1 (BRI1), BRASSINOZOLE-RESISTAMT 1 (BZR1); CONSTITUTIVE DIFFERENTIAL GROWTH 1 (CDG1); BRASSINOSTEROID-SIGNALING KINASE (BSKs); SUPPRESSOR OF BAK1(BK1); BRASSINOSTEROID-INSENSITIVE 2 (BIN2); *CYCLIN D3 (CYCD3;1); CELLULOSE SYNTHASE A (CESA*); CESA1, CESA3, CESA6; genes that encode for the proton pumps anchored in the membranes (*AHA1* and *AHA2); DWARF4* (*DWF4*), *CONSTITUTIVE PHOTOMORPHOGENIC DWARF* (*CPD*). The question mark indicates our hypothesis. Created with BioRender.com

In the osmotic gradient conditions, we observed differential regulation of the genes evaluated, some of them remained unchanged compared to the control condition as is the case of *CPD* and *DWF4* in *ttl1* and were downregulated in Col-0. Another group of genes were upregulated: *CESAs*, *CYCD3;1* and *TTL3* compared to control conditions (Figure 7). The upregulation of *CESAs*, *CYCD3;1* could, in some way, explain the root growth recovery rate of Col-0 and *ttl1* compared to the osmotic shock conditions and the absence of the swelling phenotype in the osmotic gradient. A complete transcriptome in the osmotic gradient could contribute to unraveling the gene regulatory network. The fact that the levels of *TTL3* transcript are also upregulated in Col-0 in the osmotic gradient put in evidence that in this condition another regulatory mechanism may be contributing to the phenotypes observed in the roots of *ttl1* and Col-0, like the dynamic association of TTL3 to the CSC (Kesten et al., 2022).

## Materials and Methods

### Plant Material and Growth Conditions

Columbia-0 (Col-0) was used as the wild-type genotype. The original T-DNA insertion line SALK_063943 (for *TTL1*; AT1G53300).

Seeds were sterilized in a laminar flow hood by immersion in 70% ethanol for seven minutes, then in 20% sodium hypochlorite with TWEEN®20 (Sigma, catalog number: P1379-100mL) for seven minutes, followed by five washes with sterile milli-Q water. Sterile seeds were sown on Murashige and Skoog medium (Murashige and Skoog, 1962) + 1.5% sucrose + 1.2% agar with a pH of 5.7 in square Petri dishes (12.0 cm x 12.0 cm). The plates were stratified for 48 hours in darkness at 4°C. Before seed sowing, the culture medium was autoclaved for 20 minutes at 121°C. This described medium was considered the control medium. After stratification, the sown plates were transferred to a controlled environment chamber: long-day photoperiod of 16 hours light/8 hours darkness, light intensity of 50 μEm-2s-1, temperature of 22°C, and 60% relative humidity. After seven days post-germination, seedlings were redistributed to different treatments.

### Osmotic Stress Treatment

Seedlings were grown 5 d in basal MS + 1.5% sucrose medium (−0.4 MPa) and then transferred to Petri dishes containing basal MS + 1.5% sucrose supplemented with 400mM mannitol (−1.2 MPa) or to an osmotic gradient generated with 0-400mM mannitol (−0.4 MPa to −1.2MPa). Osmotic potential was estimated by cryoscopic osmometer model OSMOMAT 030 (Gonotech, Berlin, Germany). Seedlings were grown under stress conditions for 7 days unless stated otherwise.

### Osmotic Gradient

Using a gradient maker (Supplemental Figure 1A) and a vertical container with dimensions identical to a 12.0 cm x 12.0 cm Petri dish (Supplemental Figure 1B), we created a solid growth medium with an extension of 4.5 cm. The osmotic gradient was established by mixing a 40 mL total volume of half- high concentration (20 mL) and half-low concentration (20 mL) of MS mannitol Agar solution just before pouring the separating mixture into the sandwich casting system. This mixture generates increasing osmotic potentials from −0.4 MPa to −1.2 MPa. The solution was introduced into the system by gravity, resulting in a gradient concentration between the low concentration at the upper part of the agar plate and the high one at the bottom. Based on the principle that the gradient is continuous, the position of each exact concentration of mannitol was calculated as a direct function of the distance between the initial and the final point.

Plant growth plates were assembled by sequentially placing one block of medium with 400 mM mannitol (3.0 cm), 1 block of medium with an osmotic gradient (4.5 cm), and one block of medium without mannitol (2.0 cm) (Supplemental Figure 1C). Work was carried out in a laminar flow hood to prepare the media, and materials and media were sterilized using an autoclave or ultraviolet light, depending on their characteristics. The gradient formation was assessed using the Congo Red dye (Supplemental Figure 2A) and seed germination assays (Supplemental Figure 2 B and C).

Five seedlings per genotype (7 days old) were put in each Petri dish containing the osmotic gradient. The root tips were positioned at 0 mM mannitol at the entry point of the osmotic gradient (Supplemental Figure 1C). Daily photographs were taken to measure root growth in the osmotic gradient during 25 days. The achieved osmotic potential of the roots over time was correlated using a ruler (Supplemental Figure 1 C), which links mannitol concentration to estimated osmotic potentials through a cryoscopic osmometer model OSMOMAT 030 (Gonotech, Berlin, Germany).

### Seed Germination assay

Wild-type and *ttl1* seeds (19 seeds per replicate; four replicates) were sterilized on the surface and stored at 4°C in darkness for 48 hours to induce dormancy breakage. Subsequently, the seeds were placed in each mannitol concentration strip of the osmotic gradient. After stratification, the sown plates were transferred to a controlled environment chamber: long-day photoperiod of 16 hours light/8 hours darkness, light intensity of 50 μEm-2s-1, temperature of 22°C, and 60% relative humidity. Germination percentage was calculated based on the total number of plated seeds. Pearson correlation coefficient was calculated to prove a correlation between the % of germination and the osmotic potential assayed.

### Analysis of growth rate

Growth curves were generated for Col-0 and *ttl1* in the osmotic gradient experiment. Root growth was monitored and photographed daily for 25 days. Images were captured using a Sony® Cyber-shot DSC-HX1 digital camera. Root length was measured from the hypocotyl to the tip using the free software Image J Fiji (Schindelin et al., 2012).

### Analysis of the Proximal Meristem

Roots were cleared using Hoyer’s solution (Anderson, 1954). Once cleared, the roots were mounted on a microscope slide with 40 µL of Hoyer’s solution and covered with a coverslip. Microscopy preparations were observed and photographed using the ZEISS Epifluorescence Microscope - AXIO Imager. M2 with DIC (Differential Interference Contrast) or Nomarski optics. For the proximal meristem analysis, cortex cells were counted, and their length and width were measured from the quiescent center to the first elongated cell. This point was considered the start of the elongation zone (ZE), which extended to the first root hair. The last cell of the ZE was considered the most elongated or "mature cell," and its length and width were measured (Perilli and Sabatini, 2010).

An estimate of growth parameters from cortical cell length was performed following (Cole et al., 2014) for roots grown in the osmotic gradient.

a. Root Growth Rate: (Length at day 25 - Length at the day of entry to 300 mM mannitol) / Number of days.
b. Cell Production Rate: a / Average length of the mature cell.
c. Cell Cycle Length: ((Number of cells in MP/b) * ln(2))
d. Interval Between Consecutive from TZ to EZ: 1/b

### RNA extraction

For experiments in control and osmotic shock conditions (400 mM mannitol), three biological replicates were used, each with 50-60 roots of seedlings grown for seven days under each condition. For the gradient experiment, three biological replicates were also utilized, each with 6-7 roots of seedlings grown for 25 days in the gradient, when the proximal meristem (MP) of these roots reached the position in the gradient which ranged from 300 to 400 mM mannitol. The length the roots achieved in this experiment was sufficient to extract the required amount of RNA for complementary DNA (cDNA) synthesis. RNA extraction was performed using TRIzol™ LS (Invitrogen™, catalog number: 10296028) following the manufacturer’s instructions. The RNA pellet was dissolved in 20 µL of nucleases-free DEPC-treated water (Invitrogen™, catalog number AM9915G). A 1 µL aliquot of the RNA was taken for control and subjected to DNase treatment. The RNA underwent treatment with DNase I, RNase-free (NEB®, catalog number M0303S). To 20 µL of RNA, 2 µL of 10X buffer and 0.5 µL of enzyme (5 units) were added, and the mixture was incubated at 37°C for 10 minutes in a thermoblock. Subsequently, 2 µL of 50 mM EDTA was added, and the enzyme was heat-inactivated at 75°C for 10 minutes. To assess the integrity and quantity of the extracted RNA, agarose gel electrophoresis (1%) in TAE 1X buffer was performed.

### cDNA synthesis

RNA concentration estimates were based on the visualization of bands obtained in agarose gels and their comparison with the 100 bp Plus DNA ladder (Thermo ScientificTM, catalog number: SM0321) as a reference. These estimates were used for cDNA generation. Reverse transcription reaction was carried out with the SuperScript® IV Reverse Transcriptase kit (Invitrogen™, catalog number: 18090050) using oligo d(T)20, following the manufacturer’s instructions.

### RT-qPCR

RT-qPCR was performed in the QuantStudio™ 5 Real-Time PCR System, 96-well, 0.1 mL (Applied Biosystems™, catalog number: A28138) using a PoweUp™SYBR™ Green Master Mix for qPCR (Applied Biosystems™, catalog number: A25742). For each RT-qPCR assay, three biological replicates and two technical replicates were considered. A run of RT-qPCR was conducted with biological and technical replicates of each genotype (Col-0 and *ttl1*) in the three experimental conditions (control, shock, and osmotic gradient). We tested primers of Arabidopsis AT3G18780 *ACTIN 2* and AT4G37830 *CYTOCHROME C OXIDASE RELATED* (Geng et al., 2013) gene for the housekeeping genes. As the cycle threshold was the same for both genes, we proceeded with further calculations using *CYTOCHROME C OXIDASE RELATED* as the housekeeping gene.

For fold-change calculations or the rate of change in the expression of the different evaluated genes, the Livak method (2^-ΔΔCT^) was followed (Livak and Schmittgen, 2001). It is important to note that if primer pairs used in a differential gene expression assay do not have similar and efficient amplification (between 90% and 110%), then the Pfaffl mathematical adjustment was used (Pfaffl, 2001). Primers for amplifying the housekeeping gene were included in each RT-qPCR run. Col-0 samples in control were used as the calibrator condition to normalize the ΔCT of Col-0 samples in shock and osmotic gradient and *ttl1* in control, shock, and osmotic gradient. A negative control with the absence of cDNA template was included in each experiment. The primer sequences used in the RT-qPCR assays are listed in Supplemental Table 1.

## Data analysis

Statistical analysis for differential gene expression involved conducting a two-way ANOVA test to assess genotype-by-environment interactions, utilizing INFOSTAT with a multiple testing corrected P value threshold set at 0.05 and a One-Way ANOVA utilizing Excell to analyze differential gene expression among genotypes in control conditios (Supplemental Table 3 and 4). Expression values to generate heatmap diagrams were computed using the 2^−ΔΔCT^ method, assuming ideal amplification efficiency.

Principal component analysis (PCA) of the samples and generation of heatmaps was carried out using the FactoMineR (Lê et al., 2008) and pheatmap (Kolde, 2019) packages in R Statistical Software (R Core Team, 2021). The expression values of each gene were scaled across all samples to visualize the differences.

## Author Contributions

M.S.-S. designed the research. S P-P performed root growth and RT-qPCR experiments. M.M-M performed statistical analysis. M.S.-S. wrote the manuscript. O.B. and M.M.S. critically reviewed the manuscript. All authors have read and agreed to the published version of the manuscript.

## Acknowledgments

We thanks to Gastón Quero for root growth rate estimations.

---

**Supplemental Figure 1.** System used to establish the osmotic gradient. **A.** Gradient maker. **B.** Vertical acrylic plate. **C.** Representative schematic of a Petri dish containing the osmotic gradient. Sequentially, 1 block of medium with 400 mM mannitol (3.0 cm), 1 block of medium with osmotic gradient (4.5 cm), and 1 block of medium without mannitol (2.0 cm) were placed.

**Supplemental Figure 2.** Image depicting how seedlings were planted in the osmotic gradient system. Seedlings are placed in 0 mM mannitol, and the root tips are positioned at the entry to the gradient. The ruler is shown, correlating mannitol concentrations with osmotic potentials measured using the OSMOMAT 030 osmometer model (Gonotech, Berlin, Germany).

**Supplemental Figure 3.** *TTL1* expression levels in roots of Col-0 grown in control, osmotic shock and osmotic gradient conditions. Data are presented as means ± standard deviation (SD). Three biological replicates and two technical replicates were considered, and gene expression is relative to *CYTOCHROME C OXIDASE RELATED*. Different letters indicate statistically significant differences (Tukey test P value < 0.05).

**Supplemental Table 1.** Primer’s sequences used in RT-qPCR analysis in this study.

**Supplemental Table 2:** PCA Correlation Values.

**Supplemental Table 3.** One-way ANOVA Analysis of RT-qPCR data utilizing Excell for RT-qPCR expression data in Control Conditions P value threshold set at 0.05.

**Supplemental Table 4.** ANOVA Analysis of RT-qPCR data utilizing INFOSTAT with a multiple testing corrected P value threshold set at 0.05.

